# A model of repetitive mild brain injury without symptoms – risk for Parkinson’s disease with aging?

**DOI:** 10.1101/390856

**Authors:** Praveen Kulkarni, Thomas R Morrison, Xuezhu Cai, Sade Iriah, Mary S. Lang, Laporsha Kennedy, Julia Sabrick, Lucas Neuroth, Gloria E Hoffman, Craig F Ferris

**Affiliations:** Center for Translational NeuroImaging, Northeastern University, Boston, Massachusetts; Dept of Biology, Morgan State University, Baltimore, Maryland

## Abstract

**Objectives:** To test the hypothesis that repetitive mild traumatic brain injury in early life may be a potential risk factor for Parkinson’s disease.

**Methods:** A closed-head momentum exchange model was used to produce one or three mild concussions in young adult male rats as compare to non-injured, age and weight-matched controls. Six-seven weeks post-injury, rats were studied for deficits in cognitive and motor function Changes in brain anatomy and function were evaluated through analysis of resting state functional connectivity, diffusion weighted imaging with quantitative anisotropy and immunohistochemistry for neuroinflammation.

**Results:** Head injuries occurred without skull fracture or signs of intracranial bleeding or contusion. There were no significant differences in cognitive or motors behaviors between experimental groups. Rats concussed three times showed altered diffusivity in white matter tracts, basal ganglia, central amygdala, brainstem, and cerebellum. With a single concussion, the affected areas were limited to the caudate/putamen and central amygdala. Disruption of functional connectivity was most pronounced with three concussions as the midbrain dopamine system, hippocampus and brainstem/cerebellum showed hypoconnectivity. The suprachiasmatic nucleus was isolated from all functional connections. Interestingly, rats exposed to one concussion showed *enhanced* functional connectivity (or hyperconnectivity) across brain sites, particularly between the olfactory system and the cerebellum. Immunostaining for microglia activation showed inflammation in striatum and substantia nigra with three concussions but not with one.

**Interpretation:** Neuroradiological and immunohistochemical evidence of altered brain structure and function, particularly in striatal and midbrain dopaminergic areas, persists long after mild repetitive head injury. These changes may be long lasting and serve as early biomarkers of neurodegeneration and risk for Parkinson’s disease with aging.

## Introduction

Traumatic brain injuries (TBI) are responsible for over 2.8 million emergency room visits and 50,000 deaths in the United States each year ^1^. Mild TBI is characterized as a negligible loss of consciousness with minimal neuropathology ^2, 3^ and is estimated to account for 70-90% of all TBI cases ^4, 5^. Mild TBI following a single incident is difficult to detect, and most cognitive and behavioral deficits usually resolve within weeks of the head injury, with few cases resulting in extended recovery time periods ^6–9^. However, a more pernicious, long-lasting condition may arise when the brain is exposed to repeated incidents of mild TBI (rmTBI) ^10^. Repeated mild TBI is associated with more severe and protracted cognitive, motor, and behavioral complications that may last for months and even years ^11–13^. Even after the remission of symptoms, there is accumulating evidence of persistent brain injuries ^14–18^ that carry an increased risk of dementia, Alzheimer’s disease, chronic traumatic encephalopathy, and Parkinson’s disease ^5, 19–26^.

The objective of this study was to use a momentum exchange model of repetitive head injury to produce neuropathology in a rat that would suggest risk for Parkinson’s disease (PD) with aging. It was important to establishment a specific number of head impacts that are necessary and sufficient to cause subtle or no changes in behavior in the presence of clear neuroradiological evidence of altered brain structure and function. Following disease progression over the natural life span of a rat would require the use of non-invasive magnetic resonance imaging (MRI) protocols. To these ends, we used diffusion weighted imaging (DWI) with indices of anisotropy registered to a 3D MRI rat atlas and computational analysis to identify putative changes in gray matter microarchitecture across 171 brain areas in control and experimental rats concussed one and three times. In addition, we used resting state-functional connectivity (rsFC) to evaluate changes in global functional neural circuitry. These two MRI protocols were selected due to their clinical use in diagnosing and following the progression of rmTBI after remission of symptoms ^14–18^ as well as their utility in identifying biomarkers of neurodegenerative disease ^27–31^. We used a momentum exchange model developed by the National Football League to study player concussions and designed for preclinical studies by Viano and coworkers to scale to the human experience ^32^. The velocity of head movement and energy transfer was calculated and scaled to mimic a mild concussive injury in humans producing no skull fractures, prolonged loss of consciousness, or signs of intracranial bleeding, all of which are more common in moderate and severe TBI ^33^.

## Materials and Methods

### Animals

Adult, male Sprague Dawley rats (300-400 g) were purchased from Charles River Laboratories (Wilmington, MA, USA). Animals were housed in Plexiglas cages (two per cage) and maintained in ambient temperature (22–24°C) on a 12:12 light:dark cycle (lights on at 07:00 a.m.). Food and water were provided ad libitum. All methods and procedures described below were pre-approved by the Northeastern University Institutional Animal Care and Use Committee (IACUC). The Northeastern facility is AAALAC accredited, with OLAW Assurance and is registered with the USDA. All housing, care and use followed the guidelines set by the Guide for the Care and Use of Laboratory Animals (8th Addition), the Animal Welfare Act and under the oversight of Northeastern’s IACUC which follows these doctrines.

### Momentum Exchange Model

Working with engineers at Animals Imaging Research, LLC (Holden, MA USA), we replicated the pneumatic pressure drive, 50g compactor described by Viano and colleagues and reproduced consistently, the 7.4, 9.3, and 11.2 m/s impact velocities described for mild, medium and severe rat head injury, respectively ^32^. The data reported here all came from the 7.4 m/s impact velocities as determined using high-speed video recordings. The impact piston was directed to the top of the skull, midline, in the approximate area of Bregma. All rats, control and concussed, were anesthetized with 2% isoflurane. Rats were awake and ambulatory within 5-7 min after anesthesia and concussion. Rats were observed twice daily, in the morning and early evening for the first week after concussion and weekly thereafter. Body weights were taken 2-3 times/week for the first week and then weekly. If needed, we were prepared to treat rats twice daily for three days with 0.01 – 0.05 mg/kg of buprenorphine for pain and distress, but it was deemed unnecessary based upon behavioral observations and response to handling. There were no unplanned mortalities over the course of the study.

We based our impact regimen on a rich body of data detailing the effects of acute and rmTBI in various rodent models as described ^34^. For example, Shultz et al., studied rmTBI using the lateral fluid percussion (LFP) under 2.5% isoflurane anesthesia occurring at five day intervals, with as many as five repeated injuries, and reported that the higher the number of recurrent concussions, the greater the neuropathology and cognitive and emotional deficits ^35^. Also using LFP, Aungst et al., scheduled concussions at two-day intervals, with as many as three recurrent injuries all under 2.5% isoflurane anesthesia ^36^. In still another example, Fidan et al., used the controlled cortical impact model under 2% isoflurane with one, two, or three mild concussions 24 hrs apart ^37^. Based on these and other studies, we scheduled one (n = 13) or three (n = 9) concussive head impacts under 2% isoflurane anesthesia, with a 48 hr interval between each impact. Control rats (n = 9) were exposed to isoflurane anesthesia three times with 48 hr intervals to control for the effects of anesthesia. Between 6-7 weeks after head injury, all animals were tested for cognitive and motor behavior and then imaged. At the end of the experiment 12 rats were put under deep anesthesia with 5% isoflurane and the thoracic cavity opened and the heart transcardially perfused with 4% paraformaldehyde. These brains were harvested for immunohistochemistry. The remaining rats were euthanized with a combination of carbon dioxide asphyxiation until the cessation of respiration followed by thoracotomy.

### Neuroimaging

Imaging sessions were conducted using a Bruker Biospec 7.0T/20-cm USR horizontal magnet (Bruker, Billerica, MA, USA) and a 20-G/cm magnetic field gradient insert (ID = 12 cm) capable of a 120-μs rise time. Radio frequency signals were sent and received with a quadrature volume coil built into the animal restrainer (Animal Imaging Research, Holden, Massachusetts). The design of the restraining system included a padded head support obviating the need for ear bars helping to reduce animal discomfort while minimizing motion artifact. All rats were imaged under 1-2% isoflurane while keeping a respiratory rate of 40-50/min. At the beginning of each imaging session, a high-resolution anatomical data set was collected using the RARE pulse sequence with following parameters, 35 slice of 0.7 mm thickness; field of view [FOV] 3 cm; 256 × 256; repetition time [TR] 3900 msec; effective echo time [TE] 48 msec; NEX 3; 6 min 14 sec acquisition time.

### Diffusion Weighted Imaging – Quantitative Anisotropy

DWI was acquired with a spin-echo echo-planar-imaging (EPI) pulse sequence having the following parameters: TR/TE = 500/20 ms, eight EPI segments, and 10 non-collinear gradient directions with a single b-value shell at 1000 s/mm^2^ and one image with a B-value of 0 s/mm^2^ (referred to as B_0_). Geometrical parameters were: 48 coronal slices, each 0.313 mm thick (brain volume) and with in-plane resolution of 0.313×0.313 mm^2^ (matrix size 96×96; FOV 30 mm^2^). The imaging protocol was repeated two times for signal averaging. Each DWI acquisition took 35 min and the entire MRI protocol lasted ca. 70 min. Image analysis included DWI analysis of the DW-3D-EPI images to produce the maps of fractional anisotropy (FA) and radial diffusivity (RD). DWI analysis was implemented with MATLAB and MedINRIA (1.9.0; http://www-sop.inria.fr/asclepios/software/MedINRIA/index.php) software. Because sporadic excessive breathing during DWI acquisition can lead to significant image motion artifacts that are apparent only in the slices sampled when motion occurred, each image (for each slice and each gradient direction) was screened, prior to DWI analysis, for motion artifacts; if found, acquisition points with motion artifacts were eliminated from analysis.

For statistical comparisons between rats, each brain volume was registered with the 3D rat atlas allowing voxel- and region-based statistics. All image transformations and statistical analyses were carried out using the in-house MIVA software^38^. For each rat, the B0 image was coregistered with the B0 template (using a 6-parameter rigid-body transformation). The coregistration parameters were then applied on the DWI indexed maps for the different indices of anisotropy. Normalization was performed on the maps since they provided the most detailed visualization of brain structures and allow for more accurate normalization. The normalization parameters were then applied to all DWI indexed maps. The normalized indexed map was smoothed with a 0.3-mm Gaussian kernel. To ensure that FA and RD values were not affected significantly by the pre-processing steps, the ‘nearest neighbor’ option was used following registration and normalization.

### Resting State Functional Connectivity

Preprocessing in this study was accomplished by combining Analysis of Functional NeuroImages (AFNI_17.1.12, http://afni.nimh.nih.gov/afni/), FMRIB Software library (FSL, v5.0.9, http://fsl.fmrib.ox.ac.uk/fsl/), Deformable Registration via Attribute Matching and Mutual-Saliency Weighting (DRAMMS 1.4.1, https://www.cbica.upenn.edu/sbia/software/dramms/index.html) and MATLAB (Mathworks, Natick, MA). Brain tissue masks for resting-state functional images were manually drawn using 3DSlicer (https://www.slicer.org/) and applied for skull-stripping. Motion outliers (i.e., data corrupted by extensive motion) were detected in the dataset and the corresponding time points were recorded so that they could be regressed out in a later step. Functional data were assessed for the presence of motion spikes. Any large motion spikes were identified and removed from the time-course signals. This filtering step was followed by slice timing correction from interleaved slice acquisition order. Head motion correction (six motion parameters) was carried out using the first volume as a reference image. Normalization was completed by registering functional data to the 3D MRI Rat Brain Atlas (© 2012 Ekam Solutions LLC, Boston, MA) using affine registration through DRAMMS. The 3D MRI Rat Brain Atlas containing 171 annotated brain regions was used for segmentation. Data are reported in 166 brain areas, as five regions in the brain atlas were excluded from analysis due to the large size of three brains. These brains fell slightly outside our imaging field of view and thus we did not get any signal from the extreme caudal tip of the cerebellum. Whole brains that contain all regions of interest are needed for analyses so rather than excluding the animals, we removed the brain sites across all animals. After quality assurance, band-pass filtering (0.01Hz ~ 0.1Hz) was preformed to reduce low-frequency drift effects and high-frequency physiological noise for each subject. The resulting images were further detrended and spatially smoothed (full width at half maximum = 0.8mm). Finally, regressors comprised of motion outliers, the six motion parameters, the mean white matter, and cerebrospinal fluid time series were fed into general linear models for nuisance regression to remove unwanted effects.

The region-to-region functional connectivity method was performed in this study to measure the correlations in spontaneous BOLD fluctuations. A network is comprised of nodes and edges; nodes being the brain region of interest (ROI) and edges being the connections between regions. 166 nodes were defined using the ROIs segmented from our custom MRI RAT Brain Atlas. Voxel time series data were averaged in each node based on the residual images using the nuisance regression procedure. Pearson’s correlation coefficients across all pairs of nodes (14535 pairs) were computed for each subject among all three groups to assess the interregional temporal correlations. The r-values (ranging from −1 to 1) were z-transformed using the Fisher’s Z transform to improve normality. 166 x 166 symmetric connectivity matrices were constructed with each entry representing the strength of edge. Group-level analysis was performed to look at the functional connectivity in all experimental groups. The resulting Z-score matrices from one-group t-tests were clustered using the K-nearest neighbors clustering method to identify how nodes cluster together and form resting state networks. A Z-score threshold of |Z|=2.3 was applied to remove spurious or weak node connections for visualization purposes.

### Behavioral Testing

The novel object recognition task (NOR) was used to assess episodic learning and memory ^39, 40^. The apparatus consisted of a black cube-shaped Plexi-glass box (L:60.9 W: 69.2 H:70.5 cm) with no lid, indirectly illuminated with two 40 W incandescent bulbs. Animals were placed in the empty box (15 min) for acclimation on day one. On day two, for the familiar phase (5 min), animals were placed in the box with two identical objects arranged in diagonal corners, 5 cm from each wall. After a 90 min rest period in their home cage, animals were placed back in the box for the novel phase (3 min) with one of the familiar objects and a novel object.

The Barnes Maze was used to assess spatial learning and memory as described^41–43^. The maze consists of a circular platform (121 cm in diameter, elevated 40 cm), with 18 escape holes along the perimeter at 30 cm intervals. A black, removable enclosed Plexiglas goal box was positioned under a single escape hole on the underside of the maze (L:40.0 x W:12.7 x H:7.6 cm) in the same position relative to the testing room across all trials. Between trials, the maze was rotated 45 degrees and the goal box shifted accordingly for cardinal consistency. Animals were placed inside the goal box for 1 min and then under an enclosed container at the center of the circular platform for 30 s, that was then lifted to start the trial. If animals did not find the goal box within the test period (4 min), they were gently nudged into the box and allowed to stay for 1 min, and then placed back in their home cages between trials (3 trials/day for 4 days). For both the NOR and the Barnes maze, all trials were video recorded and analyzed using manual methods by experimenters blind to treatment condition and verified with automated scoring using ANY-maze^®^ software (Stoelting, Wood Dale, IL).

A tapered balance beam (Dragonfly Inc., Ridgeley, WV) and the rota-rod were used to measure motor behavior as described ^44, 45^. The balance beam (L: 150 cm, W: 5.5 cm tapering down to 1.5 cm, elevated 120 cm) was equally divided into three sections (L:47 cm each; “wide”, “middle”, “thin” sections) that were lined with touch-sensitive sensor ledges (Width: 2 cm) that ran the length of the beam and were arranged on each side, 4 cm below the surface of the beam to count paw slips (or *foot faults*). At the start of the maze (“wide” section), was a wooden start platform, and at the end of the beam (immediately following the “thin” section) was a black enclosed Plexiglas goal box. After 2 days of training (3 trials per day), animals were tested (3 trials/day for 2 days). Prior to each trial, animals were placed inside the goal box for 1 min. Animals were then placed on a start platform and timed for traversing into the goal box, where they remained for 1 min, and were then placed back in their home cage until the next trial (30 min intertrial interval).

Following 2 days of training (3 trials/day), animals were tested over 2 days (3 trials/day) using the rota-rod by placing them on a rotating cylinder (diameter: 4 cm) that rotated at an increasing frequency starting at 1 rpm and increasing linearly at a 0.1 v/t2 acceleration rate for a total of 210 seconds ending at a max frequency of 50 rpm. Latency to fall off the rod was recorded for each animal and averaged across trials and days. For all behavior measures, GraphPad Prism version 6.0 (GraphPad Software, La Jolla, CA) was used for statistical analyses. One-sample t-tests were assessed differences from chance levels (i.e., = 50%) of exploration in the NOR task, for each experimental group individually. Comparisons between groups were assessed using one-way analysis of variance (ANOVA) or mixed ANOVAs followed by Fisher’s protected least significant difference post-hoc test.

### Iba1 Immunohistochemistry

Rats (n=3 0-hit, n=5 1-hit, n=4 3-hit) were perfused with 4% paraformaldehyde. Brains were dissected out of the skull and post-fixed for 24 hrs, then immersed in 30% sucrose in PBS until sectioning. Frozen sections (30 μm) were cut on a Microm sliding microtome and stored at −20° C in ethylene glycol/ sucrose based cryoprotectant solution until immunohistochemical processing.

Microwave-assisted immunohistochemistry was performed according to procedures described in Hoffman et al, ^46^. The primary antibody is rabbit anti-Iba1 (Wako) used at a dilution of 1:10,000. Floating sections were incubated for 48hrs at 4 degrees C on a gyrotary shaker. Secondary antibody (biotinylateed goat anti-rabbit IgG) and ABC incubations were carried out according to instructions from the supplier Vector Laboratories and modified according to Hoffman et al.^46^ Nickel-DAB was the chromagen; reagents were dissolved in 0.175M sodium acetate, pH 6.8. Hydrogen peroxide was added to a final concentration of .0025% to begin the histochemical reaction. The Ni-DAB reaction was stopped at 15 minutes by moving sections to 0.175M sodium acetate, and rinsing them several times in the same solution. Immunoreactive cells and fibers appear blue/black. Sections were mounted on gelatin coated slides, dried overnight, stained in neutral red as a counterstain for the immunoreactivity, dehydrated in graded ethanols, cleared in Histoclear and coverslipped with Histomount. Sections were examined and photographed using a Nikon microscope equipped with a digital camera assisted by IVision image software.

## Results

### Cognitive and Motor Behavior

Across days, there was a significant main effect of testing day on goal box latency in the Barnes maze test (*F*[3,81] = 9.3, *p* < 0.0001), with no significant difference between groups (*F*[2,27] = 0.38, *p* > 0.1, See Fig. 1). All groups had significantly shorter latencies to enter the goal box on testing day three (*p* < 0.0001), and four (*p* < 0.0001) when compared to the first day of testing. In addition, all groups showed shorter latencies on the last day of testing compared to the second day of testing (*p* < 0.01). In the NOR, single-sample t-tests showed that control, one, and three hit animals (*t*(11) = 6.84, *p* < 0.0001; *t*(9) = 3.86, *p* < 0.01; and *t*(7) = 4.9, *p* < 0.001, respectively) all had a significantly greater preference for the novel object, beyond chance (>50%) during the novel phase (See Fig. 1). Table 1 summarizes the results of locomotor testing showing no differences between the experimental groups.

**Figure 1.**
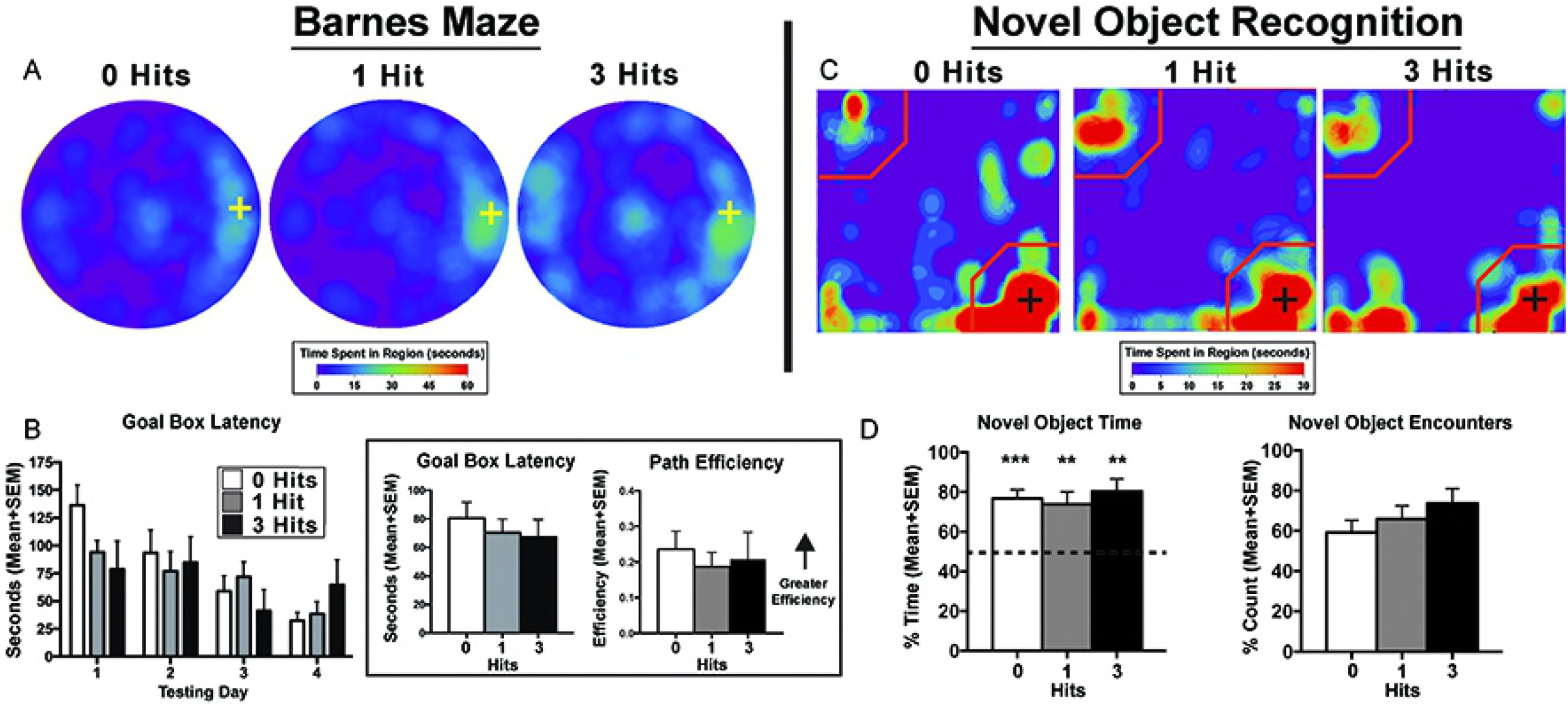
Cognition testing. (A) Barnes maze heat maps for the last day of testing. Qualitatively, on the last day of testing, animals exposed to three hits spent more time on the maze searching for the goal box, however, this effect failed to reach quantitative significance. (B) Search patterns for the Barnes maze goal box resulted in no significant differences between groups across days for all maze parameters including goal box latency and path efficiency. There was a main effect of path efficiency on the Barnes maze with all experimental groups showing relatively steady increases across days (*F*[3,81] = 3.29, *p* < 0.05). (Inset) When averaged over days, there were no significant differences between groups for goal box latency (*F*[2,27] = 0.20, *p* = 0.82) or path efficiency (*F*[2,27] = 0.22, *p* = 0.80). (C) Novel Object Recognition (NOR) – novel phase heat maps (averaged across days). Qualitative data shows an equal pattern of exploration across groups with the greatest amount of time spent in close proximity to the novel object indicated by the presence of red on the lower right corner of the maps. (D) Quantitative data reflect qualitative patterns and show that all three treatment groups spent a greater than chance amount of time (i.e., > 50%) with the novel object. Conversely, there were no differences between hit groups for the novel object recognition index (*F*[2,27] = 0.34, *p* > 0.1), total time exploring the novel object (*F*[2,27] = 0.57, *p* > 0.1), or the percentage of novel object encounters during the novel phase (*F*[2,27] = 1.16, *p* > 0.1). Crosses on heat maps indicate goal box location and novel object location on the Barnes maze and NOR, respectively. * *p* < 0.05 for single sample t-test.

### Diffusion Weighted Imaging and Quantitative Anisotropy

Measures of anisotropy at 6-7 weeks post injury were registered to the 3D MRI Rat Atlas with 171 segmented brain areas to identify possible changes in gray matter microarchitecture ^38^. The data for fractional anisotropy (FA) and radial diffusivity (RD) are shown in Fig. 2. These probability heat maps show statistical differences between the one and three hit groups as compared to controls. The effects on FA from a single concussion were limited to the dorsal striatum and central amygdala. However, rats exposed to three concussions showed significant FA changes in the olfactory system, basal ganglia, central amygdala, cerebellum, and deep cerebellar nuclei. The RD changes with one concussion were limited to dorsal and ventral striatum, medial amygdala, and trigeminal nerve. RD changes with three concussions covered motor, somatosensory, and insular cortices; dorsal and ventral striatum, globus pallidus, and the superior colliculus.

**Figure 2.**
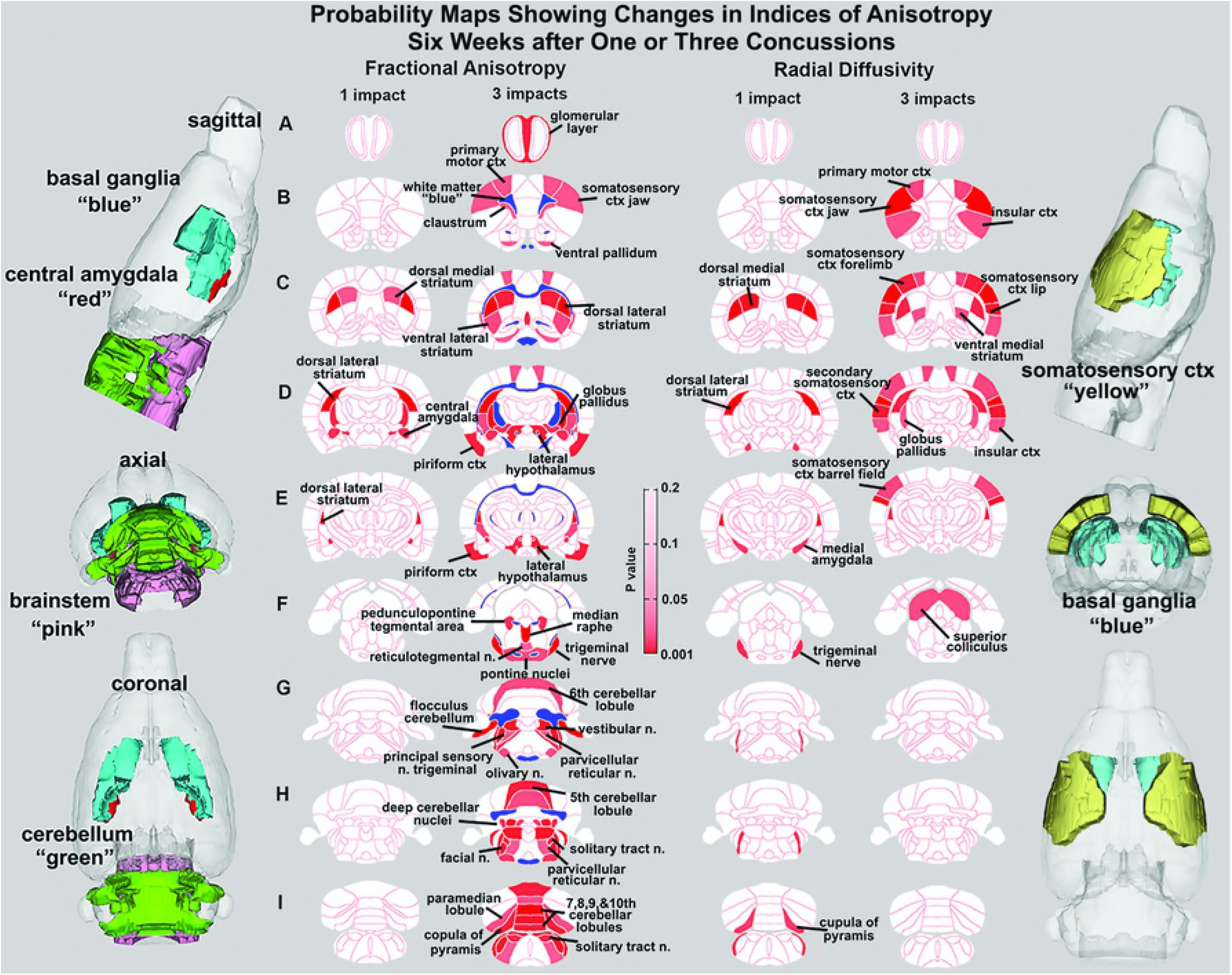
Diffusion weighted imaging. Shown are 2D probability maps with quantitative anisotropy highlighting the brain areas (pink/red) that are significantly different in FA between one (n = 13) and three (n = 9) hits versus controls (n=9) at six weeks post injury. Most of these areas are associated with the basal ganglia, cerebellum, and brainstem. The 3D representations of these areas are presented in different orthogonal directions to the left. The forebrain areas shown in sections B through D are near the impact site and include the underlying ctx and striatum (caudate/putamen), while the hindbrain areas (sections F-I) include various components of the pons and medulla oblongata that are associated with arousal (raphe, parvicellular reticular, pedunculopontine tegmental, reticulotegmental areas), sensory integration (principle sensory n, facial n, vestibular n.) and autonomic regulation (solitary tract n.). The pontine and olivary nuclei have efferent connections to the cerebellum as do the many sensory nuclei in medulla. The posterior cerebellum comprising the vermis (5^th^-10^th^ cerebellar lobules), flocculus, paramedian lobules, cupola of the pyramis, and deep cerebellar n. were all affected with three hits.

### Resting State Functional Connectivity

The delineated areas in the two correlation matrices in Fig 3 show that a single concussion favors an increase in rsFC, while repeated concussions show reduced rsFC. For example, the posterior cerebellum of the one hit group shows a much larger cluster than both the three hit and control animals. Indeed, this area has grown to include the paramedian lobule, crus 1 and 2, cerebellar lobules 7, 8, 9, and 10 plus the deep cerebellar nuclei. The rsFC between brain regions for the three experimental groups are shown in the right-hand panels of Fig 3 for the olfactory system/prefrontal cortex, suprachiasmatic n. (SCN) of the hypothalamus, and the midbrain dopaminergic system. The areas in red comprise the key nodes for each panel. For instance, the olfactory system is made up of the three layers of the olfactory bulb and the anterior olfactory nucleus. In control rats, these combined areas have significant functional connections to the marked areas of the adjacent prefrontal ctx (e.g., rostral piriform, ventral, medial and lateral orbital cortex, and the tenia tecta, highlighted in yellow). The 3D organization of these brain areas and the others is shown in the glass brains. Rats concussed once show a recruitment of functional connections six weeks post injury that includes the anterior cerebellum (3-5 lobules) and deep cerebellar nuclei (lateral and interposed). In contrast, rats exposed to three concussions have reduced connectivity that is limited only to the olfactory bulb. The SCN, the key node in the brain controlling circadian rhythms and sleep/wake cycles, has functional connections with adjacent areas of the hypothalamus in control rats that are reduced with one hit and eliminated with three hits.

**Figure 3.**
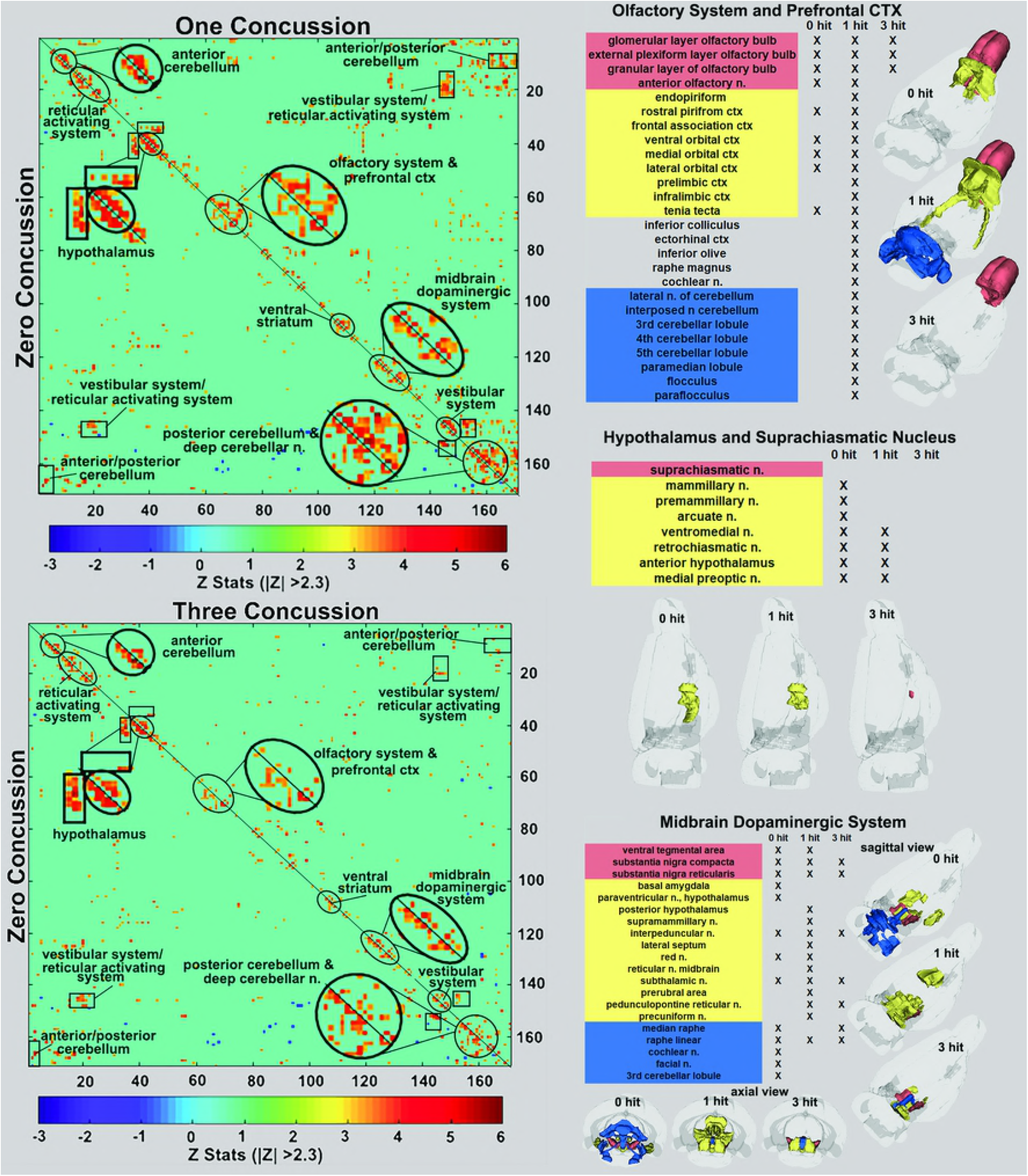
Functional connectivity correlation matrices. Shown are correlation matrices of 166 rat brain areas for rsFC comparing controls to one concussion (top left) and three concussions (bottom left). Each dark red pixel for control rats represents one of 166 brain areas that is significantly correlated with other brain areas. The brain areas with significant correlations appear as clusters because they are contiguous in their neuroanatomy and function. The diagonal line separates the control and one concussed groups. The pixels for one concussion are a mirror image of those pixels (i.e. brain areas for controls). Interesting, rats with a single concussion (n=13) showed greater rsFC within the anterior and posterior cerebellum and deep cerebellar n., olfactory system and prefrontal ctx when compared to no hit controls (n = 9).

The ventral tegmental area (VTA) as well as the substantia nigra compacta (SNc) and reticularis (SNr) make up the core nodes of the midbrain dopaminergic system. From these regions, control animals have diffuse connectivity to areas in the amygdala, hypothalamus, thalamus, medulla oblongata, and cerebellum. Following a single concussion, the functional connectivity primarily coalesces around the thalamus. Animals exposed to repeated concussions showed reduced connectivity as compared to the other groups, and had no connectivity between the SN and the VTA.

The sensitivity of the cerebellum and its efferent connections to the brain through the deep cerebellar nuclei was examined further by seeding the combined lateral, fastigial, and interposed nuclei, and mapping areas of connectivity in the one and three hit groups that were significantly different from control (see Fig. 4). In addition, the posterior cerebellum was also seeded using an aggregate of multiple areas, (6-10 lobules, cupola, crus 1 and 2, paramedian, and paraflocculus). The purpose of this seeding strategy was to identify putative afferent connections to the posterior cerebellum given its enhanced functional connectivity following a single concussion. For the one hit group, there was strong connectivity with the olfactory bulb, prelimbic ctx, tenia tecta and endopiriform ctx (section E and F). The amygdala (central, medial and basal, section D), hippocampus (CA3 dorsal and ventral, CA1 dorsal, sections D and C), motor ctx (section D) and medulla oblongata (olivary n., vestibular n. principle sensory n. trigeminal, and parvicellular reticular n., sections A and B) all showed strong connectivity to the posterior cerebellum. These connections were fewer and less significant with repeated concussions. The reorganization of functional connectivity in the cerebellum and brainstem shown in Figures 3 and 4 compliment the FA data (Fig. 2) showing alterations in water diffusion and putative gray matter microarchitecture across many of the same brain areas.

**Figure 4.**
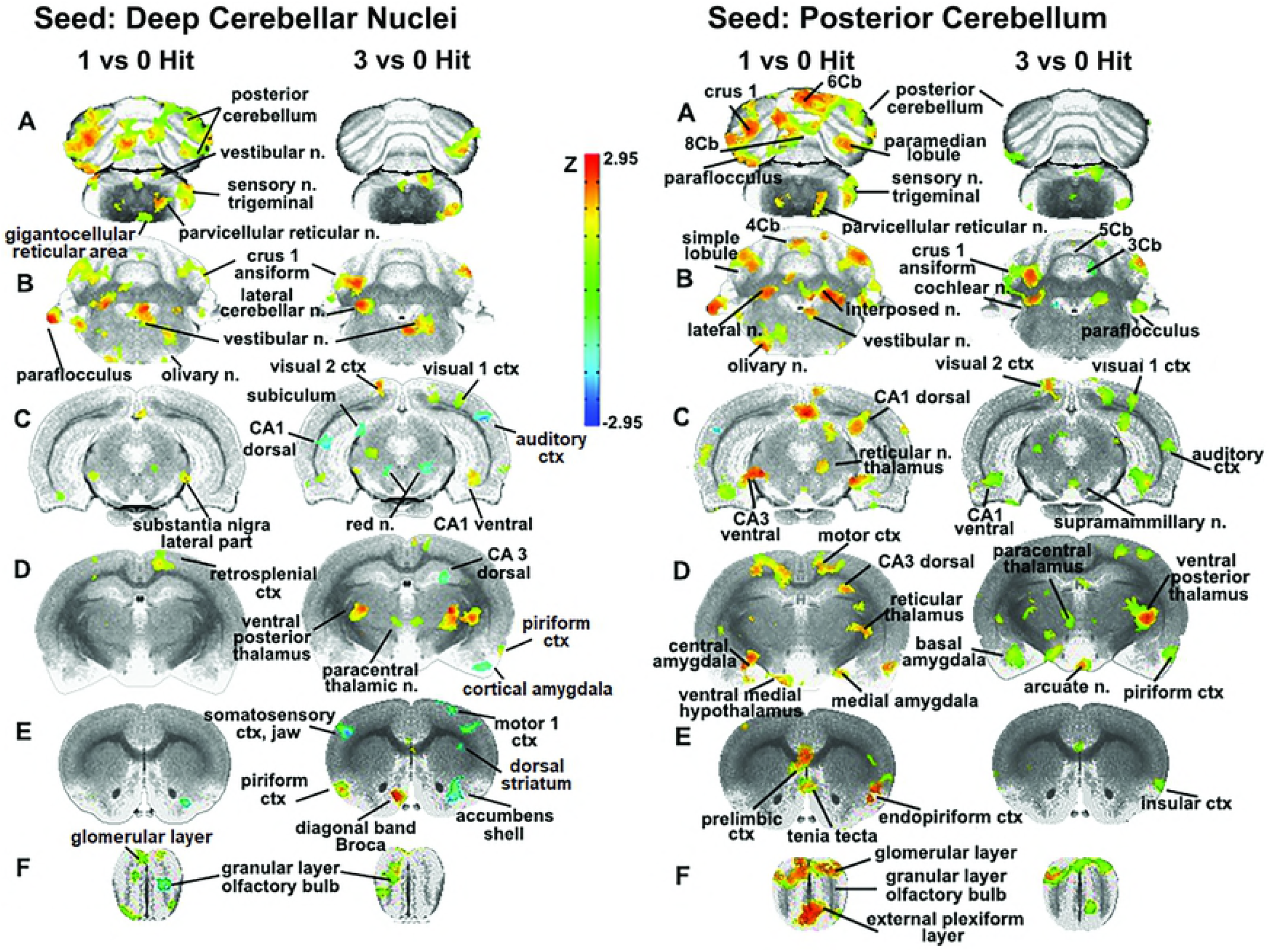
Seeding the cerebellum. The deep cerebellar nuclei (lateral, fastigial and interposed) and collective areas comprising the posterior cerebellum served as seeds point to focus on the connectivity of the brain to the cerebellum following one and three concussions (hits) as compared to 0 hit sham controls. Areas denoted in red/yellow are significantly greater than control while blue are significantly less than control. Sections A and B show increased connectivity between the deep cerebellar nuclei and the posterior cerebellum (lobules 5-9), crus 1 of the ansiform lobule, and paraflocculus in one hit rats. Additionally, the primary sensory n. of the trigeminal nerve, vestibular n., parvicellular reticular n., and the olivary n. of the underlying medulla oblongata are all part of the enhanced functional connectivity that is present in one hit rats as compared to control animals. These same cerebellar/ brain stem connections were reduced in the three hit rats (controls < three hit < one hit, sections A and B). At the level of the pons (section C) there is clear bilateral connectivity to the lateral part of the SN in one hit rats and, to a lesser degree, a unilateral connection in the three hit rats. Three hit rats show a reduced connectivity (blue) in the red n., dorsal CA1, and dorsal subiculum of the hippocampus (three hit < control) but an increased connectivity in the ventral CA1, visual 1 and 2 cortices (control < three hit). At the level of the thalamus (section D) there is only one area that differs between one hit and control animals, the retrosplenial ctx. In contrast, the three hit rats show enhanced bilateral connectivity in the ventral posterior thalamus and paracentral thalamic n., and reduced connectivity in dorsal CA3 hippocampus and piriform ctx. At the level of the striatum (section E) there is reduced connectivity in one and three hit rats in the accumbens shell (one hit ≤ three hit < control). Three hit rats show a bilateral reduction in the primary somatosensory cortex representing the jaw, a unilateral reduction in connectivity to the motor ctx, and enhanced connectivity in the piriform ctx and diagonal band of Broca.

### Iba1 Immunohistochemistry

While the differences in neuroradiology data from DWI and rsFC were consistent between one and three hit cohorts as measured by the effect size and variance, the data from IHC showing sites of neuroinflammation were more variable. In non-concussed rats, microglia were evenly distributed with a similar appearance of smallish cell bodies, with 2-5 smooth fibers. The size, appearance, and density of Iba1 immunopositive cells in the 1-hit animals were variable. Three of the one hit rats exhibited bushy microglia with a bottle brush appearance throughout the brain as shown in cerebellum (Fig 7). One of the one hit rats showed patches of microglia activation in the periaqueductal gray of the midbrain and medulla (data not shown) and another one hit rat showed patches of activation in the striatum (Fig 5). These patches of intense Iba1 staining show microglia with large cell bodies and fat, short processes. The three hit rats showed large areas of neuroinflammation that were more dispersed along the neural axis. All three hit animals showed low to high intense staining of patches in the striatum (Fig 5) while two three hit rats showed intense microglia activation in SNr and VTA (Fig 6). The cerebellum showed no patches of Iba1 staining in one or three hit rats. However, they have a bushy, bottle brush appearance indicative of activation as shown in Fig 7. These bushy microglia were found in all cerebellar lobules for one and three hit rats.

**Figure 5.**
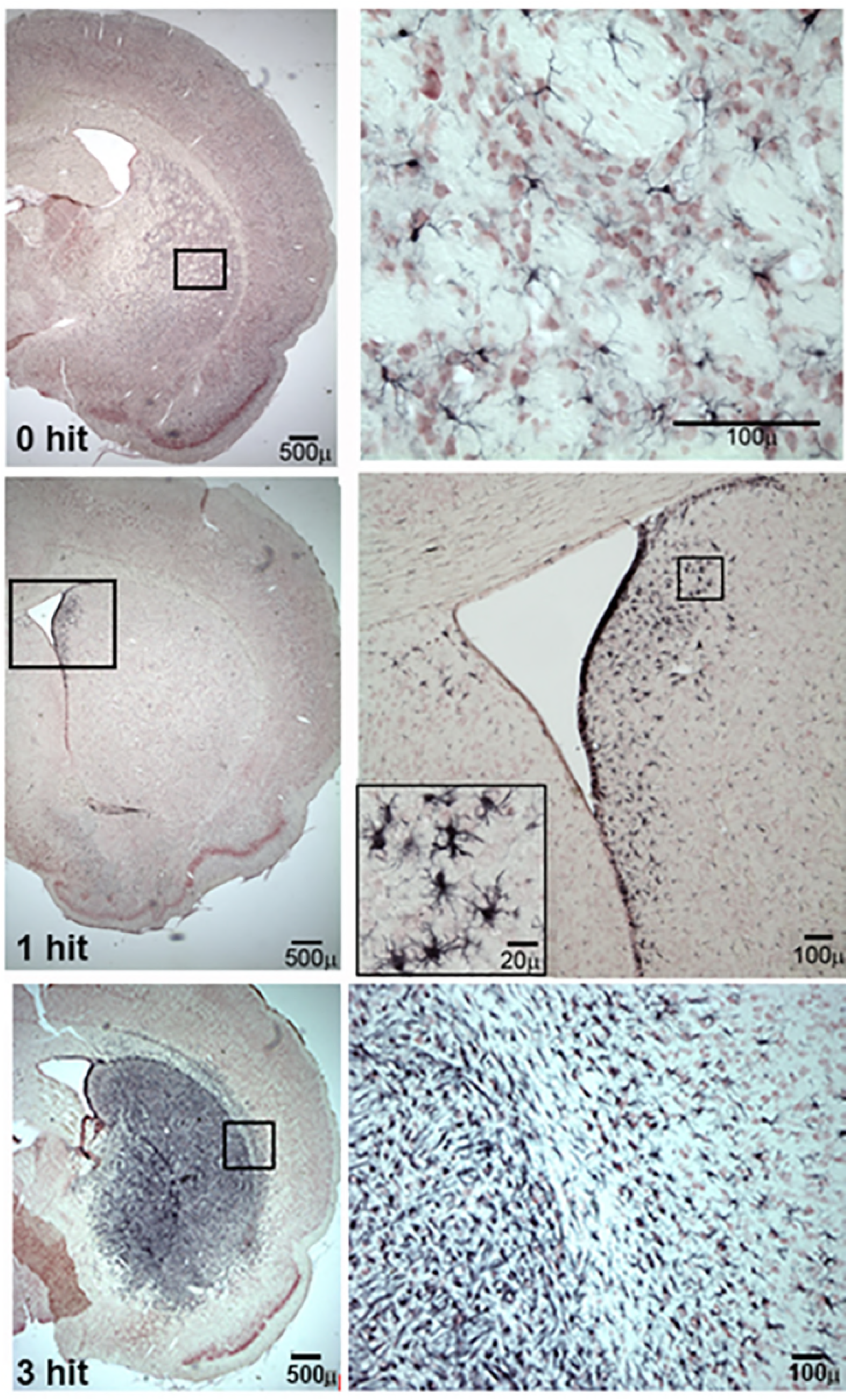
Activated Microglia in Striatum. Shown are micrographs of axial sections of the striatum (caudate/putamen) of rats concussed one and three times as compared to non-concussed rats (0 hit). Microglia immunostained with Iba1 antibody appear as multipolar black labeled cells, of which the number and density is greatly increased with three hits.

**Figure 6.**
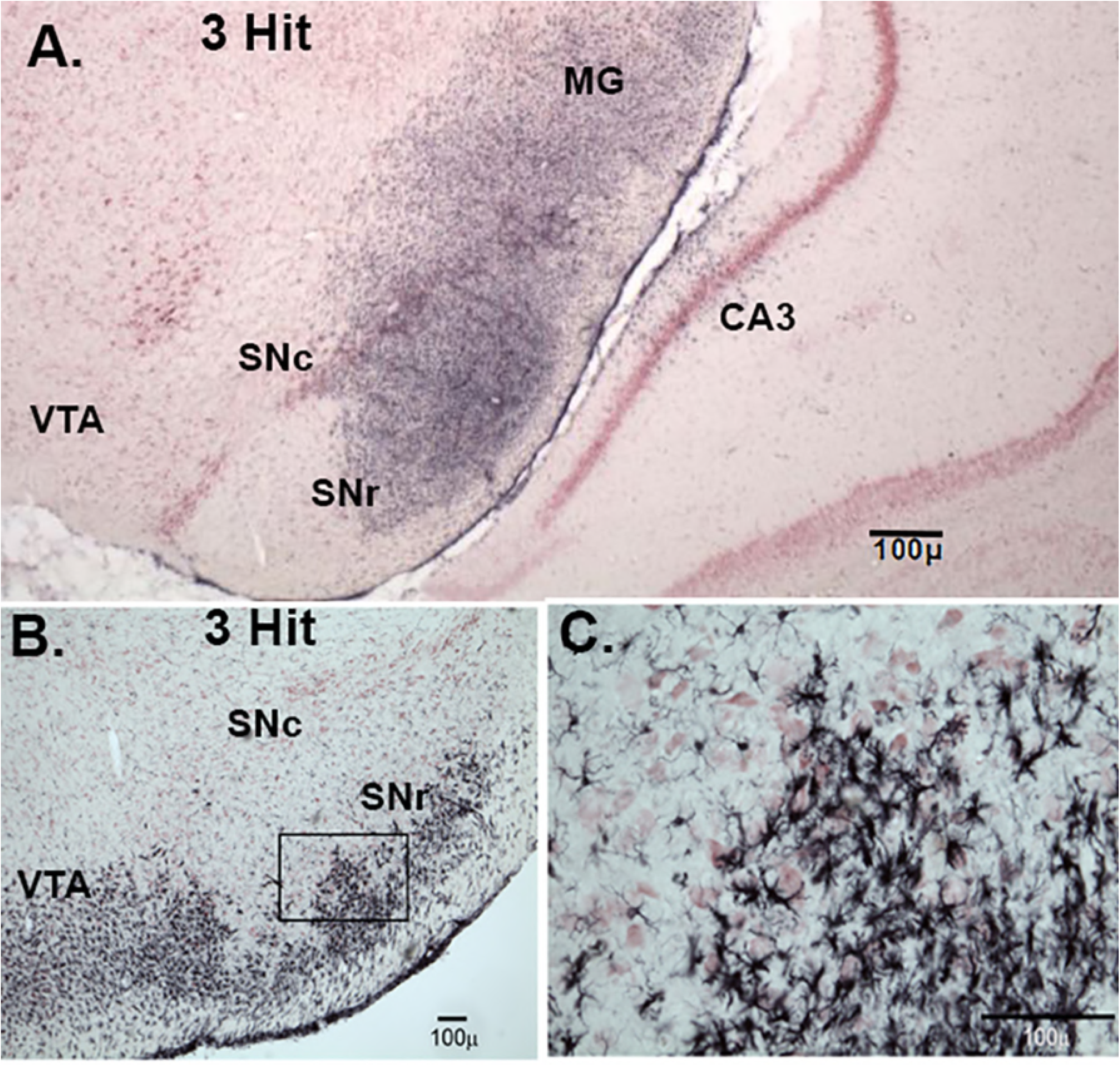
Activated Microglia in Substantia Nigra and Ventral Tegmental Area. Shown are micrographs of axial sections of substantia nigra and ventral tegmental area (VTA) from two rats concussed three times. Microglia immunostained with Iba1 antibody appear as multipolar black labeled cells.

**Figure 7.**
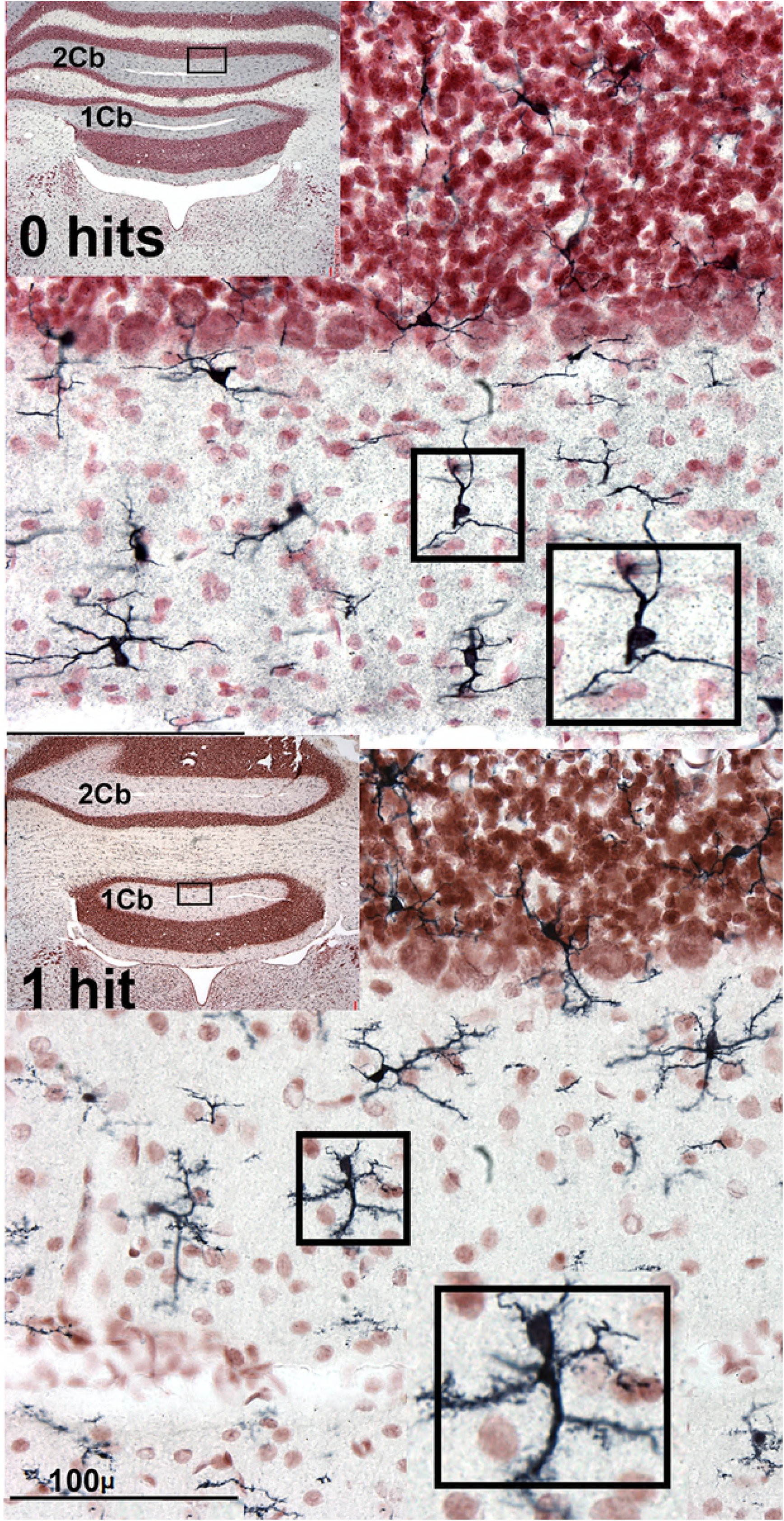
Activated Microglia in the Cerebellum. Shown are micrographs of axial sections of cerebellum from rats concussed once compared to no concussion. The higher magnification insert from each example, highlights the morphology of the microglia where the concussed rat shows ‘bushy’ microglia consistent with transition from ‘surveyor’ microglia to an activated microglia.

## Discussion

The purpose of our study was to develop a translational model of rmTBI that produced little if any overt behavioral deficits in the presence of altered brain organization and function that may pose a risk for neurodegenerative disease later in life. Indeed, we were unable to identify any changes in cognitive or motor function at six-seven weeks post injury in one or three hit rats. It is always possible that cognitive and motor deficits would have been revealed with different and more interrogative assays, but our assessment methods showed no overt problems with general health and behavior using this model of rmTBI. Nonetheless, noninvasive imaging using DWI and rsFC protocols and post mortem histology revealed significant alterations in putative gray matter architecture, functional connectivity and neuroinflammation in concussed rats with repetitive injury producing the greatest pathology.

### One vs Three Concussions

A single mild TBI caused few changes in indices of anisotropy reflecting minor alterations in central water diffusion and, accordingly, low levels of inflammation and edema as confirmed with Iba1 staining. With three concussions, there was evidence of white and gray matter injury, a loss of connectivity between various regions of the brain and neuroinflammation along the neural axis. This was not unexpected as several human and animal studies have reported repetitive injury separated by short intervals pose a greater risk than single insults or multiple head injuries separated by longer intervals ^47–53^. The vulnerability to repeated concussion with short intervals between injuries appears to involve the availability of glycolytic metabolism on demand.

Several studies have reported alterations in cerebral glucose utilization following TBI in both humans and animals. The change in metabolism following injury is triphasic with an initial period of hyperglycolysis followed by depressed glucose metabolism and finally recovery ^54–58^. In a recent study, Selwyn and coworkers looked at repeated head injury in rodents at specific time periods determined to coincide with reduced glucose uptake, and reported greater neurological damage as well as deficits in motor function, thus corroborating earlier studies that the brain needs time to recover ^56^. Concussions that occur closer together have greater cognitive and behavioral consequences, and have lasting deficits that can be present up to year later in preclinical models ^56^. Relatedly, in a repeated imaging study Qin et al., showed that, at multiple time points both during and after repeated head strikes, FA and mean diffusivity, as well as axial and radial diffusivity continue to change over time across various regions of the brain ^59^.

### Diffusion Weighted Imaging

As noted above, the changes in indices of anisotropy at six-seven weeks following a single concussion were few. The changes that occurred were localized to the central and medial amygdala, and the dorsal/ventral striatum (caudate/putamen). These areas are related to the control of emotion and dopaminergic regulation of motor function, respectively ^60–62^. Rats exposed to three concussions showed significant changes in FA within the olfactory system, basal ganglia, central amygdala, cerebellum, and deep cerebellar nuclei, while changes in RD were reflected in the motor, somatosensory cortices, basal ganglia, and superior colliculus. The resulting putative changes to gray matter microarchitecture show a distinct separation between forebrain and hindbrain (see 3D sagittal representation in Fig. 2). These results align with numerous reports showing that the cerebellum is particularly vulnerable to mild TBI ^53, 63–67^. Furthermore, in a recent study, rsFC data from human mTBI patients identified altered connectivity to the cerebellum as an important biomarker for detecting mTBI ^14^. Related to this finding, data from retired military personnel show that *decreased* metabolic activity in the cerebellum is negatively correlated with the number of blast-related mild TBIs ^63^. Given the heterogeneity of TBIs, the consistency of alterations to cerebellar function due to head injury in both humans and across animal models is uniquely distinct and suggests that the cerebellum is an important region for characterizing the progression of head injury.

### Resting State Functional Connectivity: Suprachiasmatic Nucleus

This study included a global analysis of rsFC of 166 brain regions extending from the rostral-most portion of the olfactory bulb to the caudal brainstem and cerebellum. Animals concussed once showed a combination of hyper- and hypoconnectivity across several networks, while rats concussed three times presented with only hypoconnectivity (see Fig. 3). In both one and three hit animals, the SCN had injury-dependent hypoconnectivity as compared to nonconcussed rats that showed strong connectivity between the SCN and the medial basal hypothalamus. After a single hit, the connectivity contracted to a smaller cluster, and with three hits the SCN lost all connectivity with this network.

The SCN has a critical role in circadian timing and regulation of sleep-wake cycles. Altered sleep is an important parameter in the development of our proposed animal model, as the prevalence of sleep disorders among TBI patients is approximately 50% ^68–70^. Interestingly, sleep disturbance also often appears as an early symptom in both Alzheimer’s and Parkinson’s Disease ^71–74^, and brain tissue from Alzheimer’s patients show pronounced reductions in both SCN volume and neuron density ^75, 76^. This relates to data generated using the fluid percussion model that show that TBI disrupts SCN circadian gene expression ^77^. Reduced SCN function coupled with altered sleep patterns across TBI and Alzheimer’s may suggest that a similar mechanism underlies both conditions. For example, during the preclinical stages of Alzheimer’s disease, increased amyloid-β (Aβ) accumulation coincides with the deterioration of sleep quality that, in turn, corresponds to cognitive decline and wakefulness ^78^. Similarly, in TBI, increased Aβ deposition is present acutely ^79, 80^, and years after initial head trauma ^81, 82^. A growing body of research indicates that Aβ is removed from the brain through the glymphatic system that reaches peak functionality during sleep ^83, 84^. Thus, disrupted sleep reduces the effectiveness of the glymphatic system leading to increased brain Aβ. Together these studies suggest that TBI damage to the SCN and related sleep circuits can alter circadian timing causing an accumulation of Aβ that prehaps contributes to the risk of developing Alzheimer’s disease later in life.

To our knowledge, the rsFC data is the first example showing an injury-dependent loss of connectivity in the SCN. To begin to examine the implications that this finding has for the pathogenesis of neurodegenerative disease, future studies using the momentum exchange model for rmTBI will investigate whether a loss of SCN connectivity corresponds to altered sleep patterns and/or other disturbances associated with circadian rhythms e.g., feeding, temperature, endocrine hormones.

### Resting State Functional Connectivity: Midbrain Dopamine System

The connectivity of the midbrain dopaminergic system, germane to the development of this model for the study of Parkinson’s after rmTBI, is another example of injury-dependent hypoconnectivity and reorganization of an extended neural network to a smaller cluster. The VTA and SN make up the core nodes of the midbrain dopaminergic system, and under control conditions, have a diffuse connectivity to areas in the amygdala, hypothalamus, thalamus, medulla oblongata, and cerebellum. Following a single concussion, functional connectivity primarily coalesces around the thalamus. This clustering or “small-world” effect ^18, 85, 86^ shortens the pathway length or aggregate neural connections, reducing the metabolic cost of signal transduction. Repeated concussions show reduced connectivity in these areas as compared to one hit and control groups, as well as loss of connectivity between the SN and the VTA. Both the SN and the VTA contain a high density of dopamine (DA) neurons, and the loss of functional connections with afferent brain regions due to depletion of these DA neurons is associated with Parkinson’s disease onset. The midbrain DA system and its projections to the striatum may be particularly sensitive to repeated head injury as shown here with Iba1 staining for neuroinflammation. The rmTBI-induced hypoconnectivity in the SN and VTA, along with altered DWI in the basal ganglia and persistent inflammation in the DA system observed six-seven weeks after injury may represent early risk factors for development of Parkinson’s later in life.

### Resting State Functional Connectivity: Olfactory System/Cerebellum

One of the more interesting observations from rsFC is the relationship between the olfactory system and the cerebellum. Non-concussed rats showed the olfactory bulb and anterior olfactory n. have close adjacent connections to the orbital and piriform cortices. Six-seven weeks post injury, rats concussed only once showed increased functional connections in the forebrain olfactory system and limbic ctx with hindbrain regions that include the anterior cerebellum (3-5 lobules) and deep cerebellar nuclei (lateral and interposed). In stark contrast, rats exposed to three concussions had reduced connectivity limited only to the olfactory bulb and isolated from the anterior olfactory n. The extension of connectivity across the long axis of the brain between two seemingly disparate regions with a single insult was not initially hypothesized. Both the olfactory bulb and cerebellum have numerous polysynaptic connections to much of the brain ^89–91^. However, their primary connection may be through the 5^th^ cranial nerve as the perception of many odors involves the interaction of olfactory bulbs and the trigeminal system ^92^.

BOLD imaging in response to odors that involve both systems show brain activation in the olfactory cortex, insula, thalamus, and cerebellum ^93^. Indeed, the cerebellum is consistently activated in human imaging studies that use an odor stimulus ^94^. While the pathway from the olfactory bulbs to cerebellum has not yet been defined, the circuitry appears to cross over the midline as lesions in the left cerebellum impair odor processing in the contralateral nostril ^94^. Moreover, data based on changing odor intensity suggest that the intranasal trigeminal system may be responsible for odor-induced activation of the cerebellum ^93, 95^. Disrupted olfaction is commonly found long after initial head trauma in TBI patients ^96^, and is a highly prevalent and early symptom of Parkinson’s and Alzheimer’s disease ^87, 97–99^. The differences in connectivity between the cerebellum and olfactory system with single and repeated concussions may underscore the importance of their relationship, and possibly identify novel markers for neurodegenerative disease following early head trauma.

### Resting State Functional Connectivity: Cerebellum

The sensitivity of the cerebellum and its efferent connections to the brain through the cerebellar nuclei was examined by seeding the combined dentate, fastigial, and interposed nuclei as well as the posterior cerebellum, and then mapping their areas of connectivity. The cerebellum has reciprocal interactions with much of the brain ^100^. Excitatory outputs from the cerebellar nuclei impact the motor and somatosensory cortices ^101^, thalamus ^102–104^, hypothalamus, amygdala, basal ganglia ^105–107^, and hippocampus ^108^. A single concussion increased connectivity between the cerebellar nuclei and the posterior cerebellum. The primary sensory n. of the trigeminal nerve, vestibular n., parvicellular reticular n. and olivary n. of the underlying medulla oblongata, all of which have reciprocal connections with the cerebellum ^100, 109^, and are part of the enhanced functional circuitry that we observed. The posterior cerebellum showed connectivity in the hindbrain that was similar to the cerebellar nuclei, with additional connections with the limbic ctx, amygdala, and hippocampus. The functional connectivity to these brain regions is not unexpected given the growing literature on the cerebellum’s involvement in emotion and cognition, and its reciprocal connections to these areas ^110–116^.

The altered connectivity observed after seeding the cerebellar nuclei and posterior cerebellum following three concussions showed a loss of “small worldness” with reduced connectivity with the cerebellum as well as the underlying brainstem. The motor ctx, basal ganglia, hippocampus, and amygdala showed negative connectivity as compared to nonconcussed controls. The sensitivity to these areas to rmTBI may pose a risk to the development of cognitive, emotional, and motor dysfunction associated with neurodegenerative diseases. For example, similar to our three hit model, Alzheimer’s patients show decreased functional connectivity between the hippocampus and the cerebellum ^117^, and elderly individuals with amnestic mild cognitive impairment show reduced connectivity of the hippocampus within a functional network that includes the cerebellum ^118^. In line with our three hit rsFC data, it is also notable that functional connectivity is reduced in Parkinson’s patients between the amygdala and the contralateral cerebellum, and also between the amygdala and the putamen ^119^. Combined, these studies offer support for the translatability of the momentum exchange model in regards to the etiology of Parkinson’s and Alzheimer’s disease.

### Considerations and Limitations

While behavioral assays revealed no significant differences between controls and concussed animals, it should be noted that testing occurred six-seven weeks post injury and thus, initial dysfunction may have been followed by recovery. It is generally held that the abatement of biopsychosocial deficits is accompanied by a parallel resolution of neuroradiological evidence of brain injury. While this is not the case in this study, in a recent set of experiments, Rajesh and coworkers reported that neural disruptions and structural insult in mTBI may persist up to 10 years following injury in subjects with normal cognitive function ^17^. Hypoconnectivity in the forebrain proposed to be responsible for initial cognitive deficits persisted for years after injury suggesting the brain may compensate for disrupted function through reorganization. A time lapse of six-seven weeks in a rat’s life is comparable to 4-5 years in humans, and thus the continued presence of injury after rmTBI suggests that the hyper- and hypoconnectivity and neuroinflammation observed in these studies may persist over the life of the rat.

The site of impact was limited to the rostral cranium at the level of bregma directly affecting the underlying motor ctx. Striking this specific site may have produced a unique mechanical force responsible for the observed global changes. However, we believe that the neurological effects of TBI are more generalized and agnostic to the site of impact. While concussions can occur on any part of the head, the general neuropathology is reasonably similar between cases. The cerebellum has been recognized as being particularly vulnerable to mTBI ^14, 63, 64, 120^ and neuroradiological evidence of cerebellar dysfunction has been promoted as a diagnostic biomarker of TBI ^14^. Previously, we addressed whether general markers of dysfunction reliably occur after TBI between subjects and found that concussive injuries to the forebrain or hindbrain of rats results in a similar pattern of neuropathology in the amygdala, hippocampus, and thalamus ^38^.

The rsFC data were collected under low dose isofluorane anesthesia to minimize motion artifact and physiological stress ^121^. Nonetheless numerous studies comparing the anesthetized and conscious states show similar rsFC data ^122, 123^. However, the absence of anti-correlation data in these studies may be attributable to the anesthesia ^124^.

## Summary

Recent clinical studies report mild TBI early in life is a significant risk factor for future dementia and Parkinson’s disease^26, 125–127^. The momentum exchange model developed by the National Football League to study player concussions was adopted for use on rats to produce mild concussions with no neuroradiological evidence of brain contusions or changes in cognitive or motor behavior. Nonetheless, six-seven weeks post injury there are significant changes in brain morphology and function as determined by MRI. The midbrain dopaminergic system and striatum are particularly vulnerable to rmTBI. Post mortem analysis for activated microglia show neuroinflammation in striatum, SNc and VTA. The imaging data fit the findings reported in the clinic years after head injury in humans. This momentum exchange model combined with non-invasive MRI to follow disease progression parallels the human experience and provides an opportunity to investigate the efficacy of early intervention strategies impacting dementia and Parkinson’s with aging.

## Acknowledgements

Program Consortium BUILD Award (UL1MD009605/RL5MD009590/TL4MD009635)

